# Data-driven design of LNA-blockers for efficient contaminant removal in Ribo-seq libraries

**DOI:** 10.1101/2025.09.11.675547

**Authors:** Dario A. Ricciardi, Franziska E. Peter, Maik Böhmer

**Affiliations:** Institute of Molecular Biosciences, Plant Physiology, University of Frankfurt, Max-von-Laue Str. 9, 60438 Frankfurt am Main, Germany

**Keywords:** Ribo-Seq, rRNA, depletion, LNA, sequencing, translation, Arabidopsis thaliana

## Abstract

Ribo-Seq libraries often contain a high amount of non-coding RNA fragments, which can significantly reduce the information output of these experiments. Contaminants can comprise up to 90% of a Ribo-Seq library, showing high sequence variability and diverse fragmentation, which hinders the effectiveness of rRNA depletion kits with fixed target sequences. We developed a workflow to identify experiment-specific contaminants from a small-scale, preliminary sequencing run. This enables the design of locked nucleic acid (LNA) oligonucleotides that target the contaminating fragments, thereby preventing their amplification during library preparation. This process requires only a single pipetting step and no additional purification. In a proof-of-concept experiment, just five LNAs reduced contaminating fragments by over 30 %, doubling the amount of useful sequencing data from Ribo-Seq experiments.

We offer a script to identify and visualize contaminants and optimized target sequences, along with guidelines for designing custom LNA sets and a collection of predesigned LNAs for *Arabidopsis thaliana* across various common growth conditions, serving as a foundation for a public LNA repository.

**Significance Statement:** Ribo-Seq libraries often contain abundant non-coding RNA contaminants, which, because of their high sequence variability and diverse fragmentation, are challenging to remove. We present a computational pipeline that identifies experiment-specific target sequences and allows for their efficient depletion using custom LNA probes in a single pipetting step, thereby increasing sequencing yield and reducing costs. A public LNA repository will support sharing validated targets within the research community.

## Introduction

Ribosome profiling (Ribo-Seq) is a relatively new technique that enables the quantification of regulatory events at the translational level, similar to RNA-Seq at the transcriptional level (Ingolia *et al*., 2009). For a Ribo-Seq analysis, ribosomes are extracted from tissue in their native state, still bound to actively translated mRNA transcripts. Exposed mRNA not protected by the ribosome is degraded by nuclease digestion, while the ribosome-protected fragments (RPFs), or ribosomal footprints, are purified and sequenced. RPFs are then mapped back to the reference genome to reveal the exact distribution of translating ribosomes, producing a snapshot of the translatome. Especially in conjunction with transcriptomic data, this information can be used to infer translation efficiency, slow/pause sites, and alternative reading frames, making Ribo-Seq an essential technique for understanding translational regulation (Brar and Weissman, 2015).

The mammalian and microbiological fields have already recognized the significant potential of regulatory information from exploring the previously overlooked area of translational control of biological processes (Tierney *et al*., 2025). There are well-established Ribo-Seq protocols for mammalian cell systems and microbes. In contrast, protocols for plants and other less common model organisms are still developing, and commercial kits are limited.

However, plants may show an even greater level of translational regulation. They have unique mechanisms of controlling translation, such as using upstream open reading frames (uORFs) and reprogramming translation in response to stress. These methods are less common in yeast and mammals, where translational regulation is usually limited to specific stress responses (Browning and Bailey-Serres, 2015; Izquierdo *et al*., 2018). Especially during stress conditions like hypoxia, heat, or dehydration, Arabidopsis exhibits a general reduction in polysome-bound mRNAs, with many mRNAs stored in non-polysomal RNPs. This targeted repression of translation is reversible and helps plants conserve energy while prioritizing the translation of stress-related mRNAs (Juntawong *et al*., 2014; Branco-Price *et al*., 2008).

The success rate of a Ribo-Seq experiment depends on the yield and quality of the RPFs and the resulting sequencing library. The processing steps required to obtain RPFs often result in Ribo-Seq datasets containing significant proportions of contaminating fragments of non-coding origin.

These contaminants predominantly originate from ribosomal RNA that is cleaved at specific regions accessible to the nuclease used for footprinting (Supporting Figure S1). This selective cleavage results in a deterministic distribution of a few highly abundant contaminant fragments that co-purify in the same size range as RPFs (Supporting Figures S2).

Established rRNA mitigation strategies that rely on probes with pre-defined target sequences against ribosomal RNA (rRNA depletion) or mRNA (mRNA enrichment) underperform when applied to Ribo-Seq samples due to the fragmentation of their targets (Zinshteyn *et al*., 2020). This poses a challenge for commercially designed kits because the contaminant profile is influenced by the organism, the growth and experiment conditions, and the nuclease used for footprinting, leading to many different combinations of possible contaminants (Gerashchenko and Gladyshev, 2017).

The specificity and therefore the efficiency of the depletion have been improved significantly by using custom-tailored probes, coupled with affinity purification (Ingolia *et al*., 2012; Brar *et al*., 2012; Gerashchenko *et al*., 2012; Alkan *et al*., 2021; Ting *et al*., 2024), nuclease digestion (Chung *et al*., 2015; Baldwin *et al*., 2020; Gu *et al*., 2021), or Cas 9 cleavage (Gu *et al*., 2016; Han *et al*., 2020). However, these methods need more hands-on time, incubation steps, and especially purification steps, which could cause sample loss and biases.

Oligonucleotides containing locked nucleic acid (LNA), which are chemically modified nucleotides with a fixed conformation that enhances binding affinity, can effectively bind contaminant fragments during library preparation and sterically prevent reverse transcription or amplification, eliminating the need for additional cleanup steps (Prout *et al*., 2023).

We present a workflow that efficiently removes contaminants from Ribo-Seq libraries in a single pipetting step using LNA probes. This method utilizes software to identify experiment-specific contaminants from an initial sequencing run. Additionally, we offer contaminant profiles for common growth conditions of *Arabidopsis thaliana*.

## Results

To enable targeted removal of highly abundant contaminant fragments from Ribo-Seq libraries using locked nucleic acid (LNA) oligonucleotides, it is first essential to identify the various RNA species produced by ribonuclease activity during the footprinting step. The most effective method is to perform a preliminary sequencing experiment with a smaller sample size and lower sequencing depth under the planned experimental conditions. This allows for the characterization of contaminant fragment profiles specific to the organism, growth conditions, treatment, and the ribonuclease used for mRNA digestion. As a simpler alternative, we provide indicative contaminant distributions for *Arabidopsis thaliana* grown under standard conditions and processed with RNase I for footprinting. Even more simply, we present five LNAs that are highly abundant across all tested growth conditions and have been tested for their efficiency in contaminant removal.

### Contaminant Identification Script

To identify highly abundant contaminant RNAs, we developed an R script that utilizes pre-aligned reads from a small-scale sequencing run. Reads are assigned to genetic features, quantified, and grouped based on sequence similarity. The shortest common sequence in each group is reported as the optimal LNA target sequence (Supporting Figure S3). The script takes only a few minutes to analyze a small dataset (10–20 million total reads) on modern hardware, thanks to an optimized sequence aggregation process. First, only the top 10,000 sequences are used as search patterns, which are then used to find similar sequences across the rest of the file. Second, already matched sequences are removed from the set of search patterns, significantly reducing the number of matching operations required.

Matching parameters can be customized. However, reducing the matching depth speeds up computation but results in less optimized target sequences. Higher matching depths often produce groups with shorter common target sequences. The overall distribution of contaminants is only slightly affected by the matching depth. The default grouping mode (“shortest”) summarizes reads based on their shortest common sequence; however, there is also a mode that allows for mismatches or shifts at either end (“fuzzy”). The “fuzziness” parameter defaults to three nucleotides and effectively trims the sequence internally at each end to find similar sequences. The third mode (“sequence”) only summarizes exact sequence matches, which effectively creates an abundance table with each unique sequence in the dataset.

The mapping function requires BAM files with exactly one alignment per read. For multimapping reads, which comprise the majority of contaminants, the output should be the top-scoring alignment. In this work, we used the STAR mapper (Dobin *et al*., 2013a) with the settings -- outSAMmultNmax 1 and --outFilterMultimapNmax 20 to allow for multimapping reads while writing only a single alignment for each, ensuring accurate quantification. The script works with any organism, as long as a genetic annotation file is provided in either GTF or GFF format. Running the script produces three main outputs: A comma-separated table containing the most abundant fragments, grouped by their shortest common sequence with all associated genomic features, a heatmap visualization of the distribution of contaminating fragments across all samples, and a figure of merit to help the user gauge the expected depletion performance in relation to the number of LNA targets (Figure 1).

**Figure 1:**
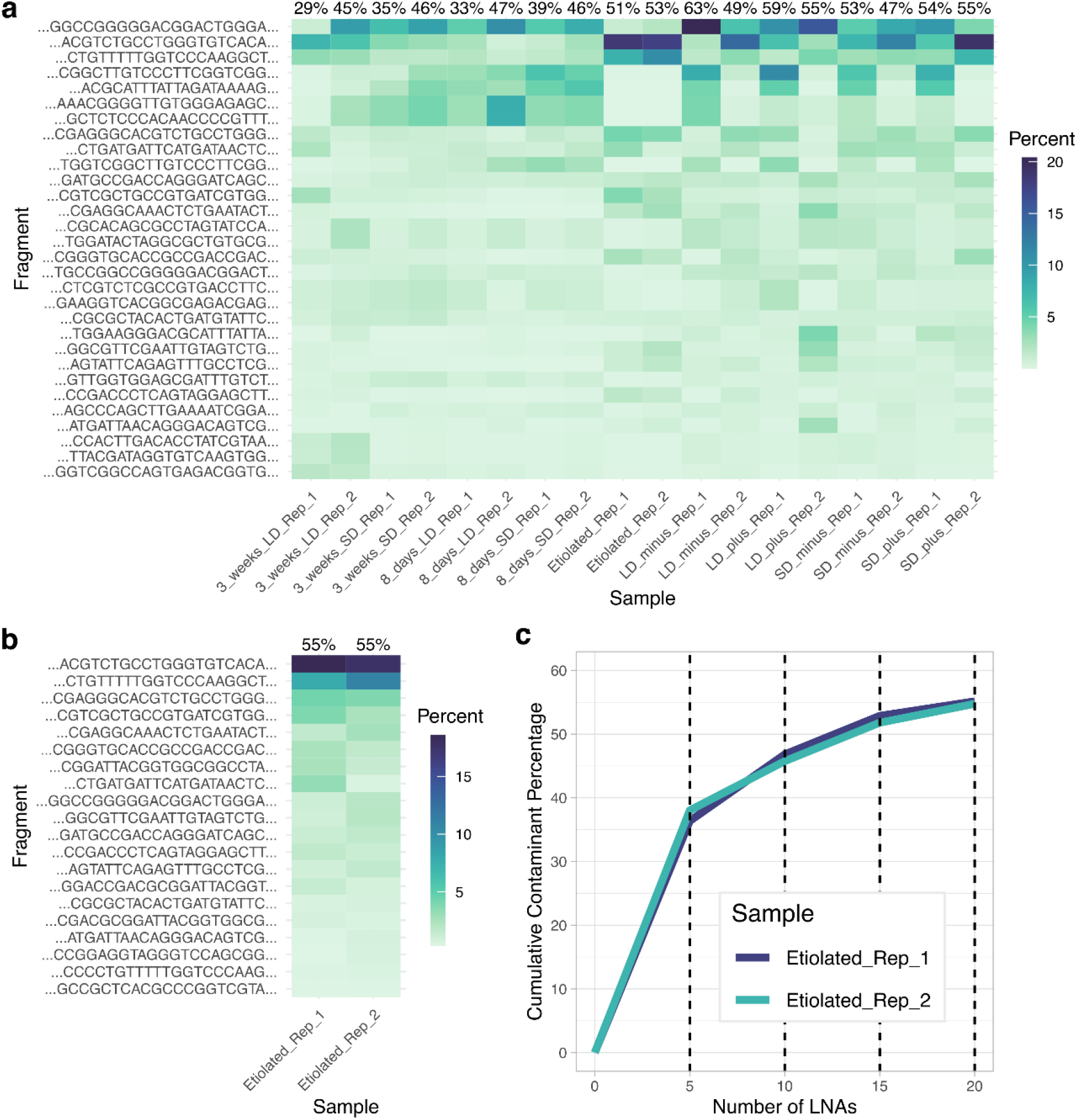
Visualization of contaminant profiles across different growth conditions. Arabidopsis seedlings were grown for 8 days or 3 weeks in soil or hydroponically for 8 days in ½ MS medium with and without sucrose (plus/minus). Light conditions were either short-day (SD) or long-day (LD). One set of plants was grown hydroponically for 8 days in ½ MS medium without sucrose in total darkness (etiolated). RNase If was used for the footprinting reaction. **a**: Heatmap visualization of the 30 most abundant sequences after analysis with our script. Sorted by average abundance across all conditions. Each row represents a group of similar sequences of varying lengths, sharing a shortest common sequence, which is displayed as the y-axis label. Tile color represents the percentage of sequences in that group in relation to the total number of sequences in the sample. The cumulative percentage of the grouped sequences for each sample is indicated at the top of each column. **b**: Heatmap visualization of the 20 most abundant sequences in a single condition (etiolated seedlings). **c**: Figure of merit for estimating the optimal number of LNA targets for etiolated seedlings. Percentages of each target are added by walking down the sequences from top to bottom. Results are binned every 5 targets.

### Contaminant Profiles of Common Growth Conditions

To assess whether different growth conditions would influence the occurrence and quantity of contaminant fragments during Ribo-Seq library preparation, *Arabidopsis* plants were grown for either eight days or three weeks under varying conditions. The plants were harvested and processed for Ribo-Seq by generating and purifying ribosomal footprints, which were subsequently converted to sequencing libraries. The sequenced reads were aligned and analyzed with our contaminant identification script.

All tested conditions were visualized in a unified heatmap to directly compare the distribution of contaminant fragments across samples (Figure 1a). Soil-grown plants, regardless of age and light conditions, exhibited similar profiles, with a high abundance of contaminants concentrated in the top seven groups. One of the samples (3 weeks, long-day, replicate 1) exhibited a significantly different distribution of contaminants, with only two groups aligning with the rest of the soil-grown plants. A single sample with such a diverging profile could be considered an outlier; however, the distributions of contaminants among the plate-grown plants showed similar heterogeneity. While most contaminants were grouped at the top, indicating common sequences with all other tested conditions, every other replicate had a contrasting distribution that did not align with the other replicates from those conditions. Indeed, we observed that while age and growth conditions had a slight effect on the occurrence and distribution of contaminant sequences, a sizable impact could be traced back to the batches in which the samples were processed, suggesting a strong influence from the digestion conditions during sample preparation.

One condition that stood out was the growth in the dark, which resulted in etiolated seedlings. Etiolation drastically reshapes the translation machinery, especially in chloroplasts, and leads to an altered abundance of ribosomes, which contribute most of the contaminating sequences. The contamination profile of etiolated seedlings was antithetical to all other conditions. It showed a high concentration of sequences in a single group originating from the 3’ end of the 5.8S rRNA.

Including all growth conditions in a single plot and sorting by average abundance provided a good comparison across multiple growth conditions; however, most researchers usually test only a few conditions relevant to their specific experiment setup. Inspection of a single condition yielded a clear ranking of contaminants, with a notable decline after a limited number of entries (Figure 1b). The script provides a figure of merit based on the ranking, which helps the user determine expected depletion efficiencies and decide how large the set of LNAs should be for the experiment (Figure 1c). The figure of merit showed diminishing returns as the number of LNA targets increased, leaving it up to the user to determine the most cost-effective point. We predicted a 15– 40% reduction in contaminants using a set of five LNAs targeting the most abundant sequences, based on a figure of merit across all tested conditions. Then, we aimed to compare the projected reduction with the actual depletion efficiency.

### LNA Design and Application

After identifying the top 5 targeted sequences across all conditions, LNA probes were designed for optimal binding efficiency according to Prout *et al*. (2023). In summary, LNA oligonucleotides consisted of alternating DNA and LNA nucleotides, starting with a DNA nucleotide (Figure *2*a). We recommend a minimum length of 14 nucleotides in total, mainly due to sequence specificity concerns but also for practical reasons. Ribo-Seq libraries typically do not contain sequences shorter than 20 nucleotides, and aligning such short reads is unreliable. Additionally, we phosphorylated the 3’-end to prevent the LNA oligonucleotides from acting as primers themselves and producing a truncated amplicon that could interfere with downstream analysis. The properly designed set of LNAs was ordered in medium quantities (200 nM synthesis scale) with HPLC-grade purity. We suspect less pure oligonucleotides will work similarly well.

**Figure 2:**
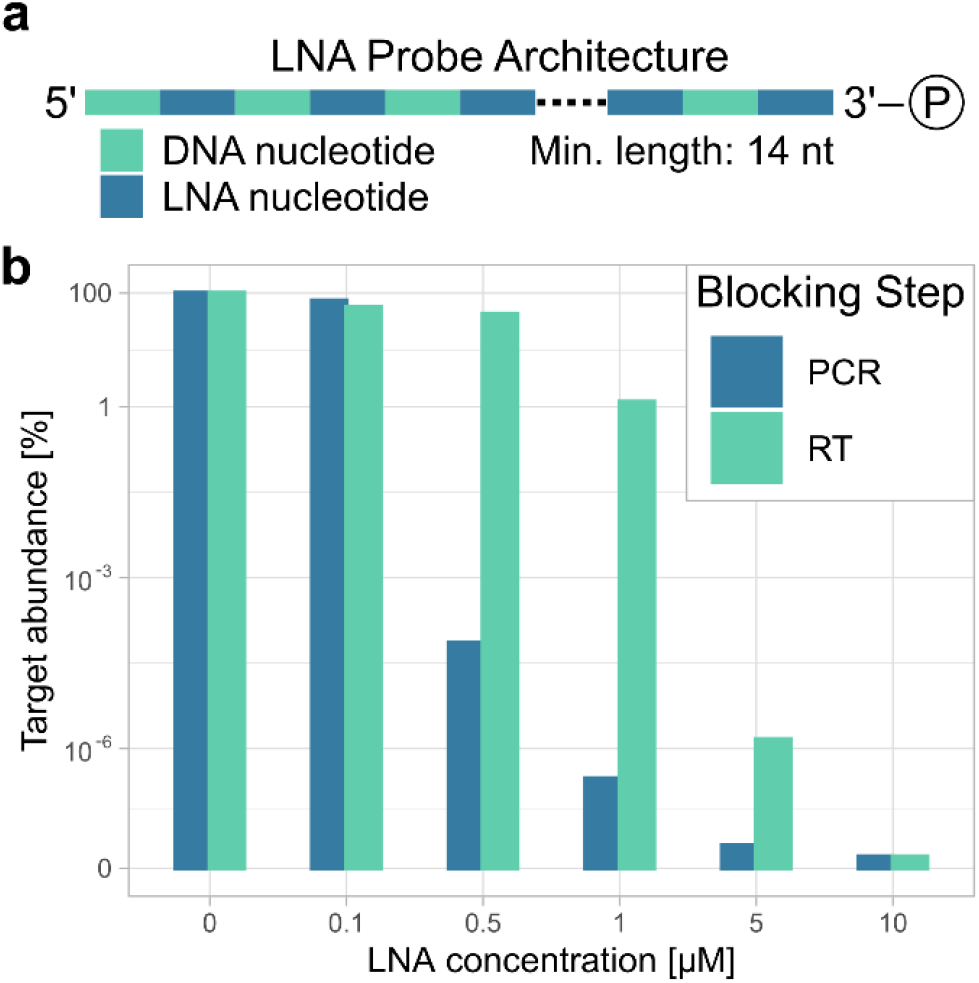
LNA design and application considerations. **a**: Optimal architecture of LNA probes. Alternating DNA and LNA nucleotides, beginning with a DNA nucleotide. The dotted line represents variable length. Phosphate residue at 3’-end. **b**: Blocking performance of a single LNA against its target (fragment of 5.8S rRNA) at various concentrations. Comparison between blocking at the reverse transcription step and during PCR amplification. Target abundance percentages are based on qPCR cycle difference in relation to the non-depleted sample (0 µM).

LNAs can be added before reverse transcription or prior to PCR amplification of the library. To determine the optimal step, we conducted preliminary quantification of the blocking performance using primers for the top contaminant and adding increasing concentrations of the appropriate LNA probe at either the reverse transcription step or at the amplification step (Figure *2*b). RT-qPCR was used to quantify the abundance of the target sequence in the Ribo-Seq library.

In short, total RNA was extracted from Arabidopsis seedlings, reverse-transcribed into cDNA, and subsequently amplified in a qPCR cycler. 10 µM of the tested LNA completely blocked target amplification in both the reverse transcription and amplification steps, with no product detectable. At lower concentrations, the tested LNA demonstrated superior blocking efficiency during PCR amplification compared to the reverse transcription reaction. The addition of 0.5 µM LNA to the PCR reaction resulted in a more than 1000-fold reduction in product abundance. Application of the same concentration during the reverse transcription step only reduced the product by a few percent. Adding the LNA probes during amplification provides the additional advantage of strand agnosticism. Blocking either the forward or reverse product effectively diminishes amplification efficiency, simplifying LNA probe design by eliminating the need to match the template sequence during reverse transcription.

### LNA Depletion

After identifying the top contaminants and optimizing the concentration and application of the LNA blockers, we designed a set of five LNAs targeting the top five contaminant fragments across all tested conditions as a ready-to-use alternative to custom probe sets (Table 1). Ribosome-protected-fragments were purified from hydroponically grown, untreated wild-type Arabidopsis seedlings. Pooled extracts were split and converted into four identical sequencing libraries. LNA blockers were added to two libraries during the amplification stage at a concentration of 1 µM each. The resulting reads were aligned and analyzed with our script. Targeted depletion of the top five contaminants yielded a reduction of over 30 % of the total identified contaminants (Figure 3a). Sequences that were targeted by the blockers were eliminated efficiently, going from 8–12 % of total reads to less than 1 %.

**Table 1:**
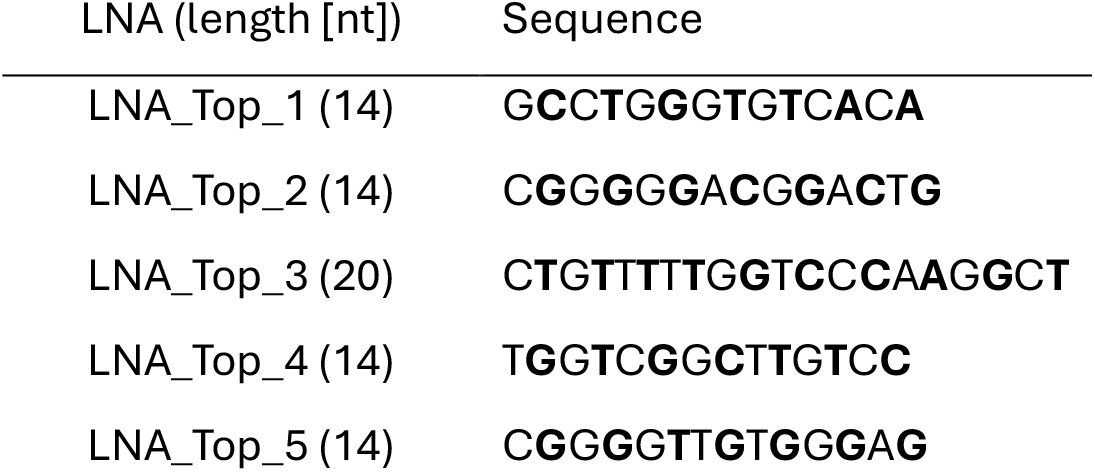
Locked nucleic acid (LNA) oligonucleotides used for the depletion of the most abundant contaminants across all Ribo-Seq libraries. LNA nucleotides are denoted in bold letters.

**Figure 3:**
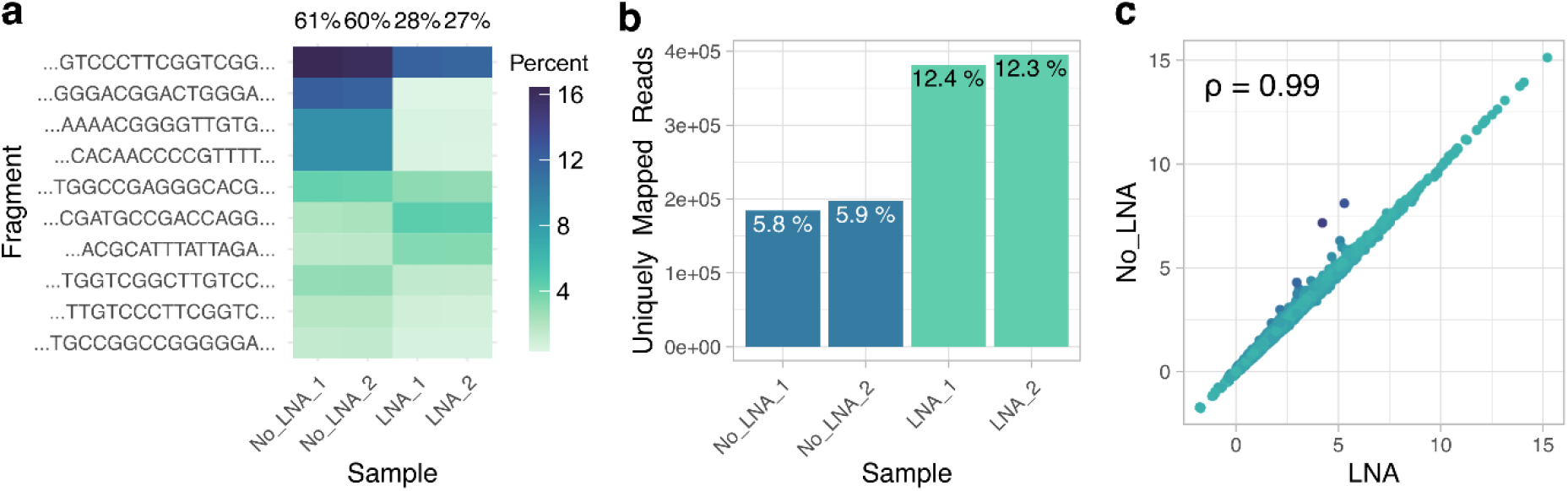
Depletion performance of five LNAs targeting the most abundant contaminants. **a**: Heatmap visualization of undepleted samples and samples that were supplemented with an LNA mixture during library amplification. The LNA mix contained five probes at 1 µM each, targeting the most abundant contaminants identified across all tested growth conditions. The shortest common sequence of each contaminant group is indicated on the y-axis. Tile color represents the percentage of that group in relation to the total reads in the sample. Cumulative percentages for all displayed contaminants are summarized at the top of each column. **b**: Uniquely mapping reads in undepleted and LNA-depleted samples. Uniquely mapping reads mainly originate from coding sequences and are the desired readout of Ribo-Seq experiments. Percentages in relation to total read count per sample are indicated at the top of each bar. **c**: Correlation of averaged read counts between undepleted and depleted samples. Counts were regularized log-transformed using the “rlog” function from the DESeq2 R package. Pearson correlation coefficient is denoted in the top left corner.

The relative abundance of non-targeted contaminating fragments increased due to the depletion of the highly abundant targeted sequences. More importantly, the yield of RPFs of coding origin, which was the desired outcome of the depletion, significantly increased (Figure 3b). We observed a near-exact doubling of uniquely mapping RPF reads. To verify that the depletion did not adversely affect read counts from transcripts of interest, we correlated the rlog-transformed average counts from undepleted libraries with those from LNA-depleted libraries, considering only uniquely mapping reads (Figure 3c). We saw a near-perfect correlation between the two datasets with a correlation coefficient of 0.99 (Pearson). Two outliers were detected, which seemed to be affected by the depletion. Individual inspection of the aligned reads in the Integrative Genome Viewer, however, revealed misaligned reads from a multimapping origin, which were intended to be targeted by the depletion. The actual uniquely mapping reads were not affected by these transcripts.

## Discussion

Ribo-Seq is inherently a complex technique that can pose significant challenges for researchers aiming to decipher translational regulation. To address these difficulties, we developed a user-friendly software tool that obviates the need for extensive bioinformatics expertise, provides comprehensive guidance on the design and application of effective LNA probes, and initiates a shared library of validated LNA sequences. This workflow successfully reduces sequencing costs by enhancing the yield of informative ribosome-protected fragments (RPFs). Additionally, by consolidating contaminant depletion into a single streamlined step, we have substantially improved the method’s accessibility, particularly for researchers who want to start performing Ribo-Seq experiments.

The custom LNA-based depletion can also be leveraged in established workflows, where it provides greater specificity, flexibility, and cost-effectiveness. Particularly, the latter may be an advantage over commercial offerings, especially in multiple experiments. Though LNA probes are more expensive than pure DNA oligonucleotides, they often amortize after the first use, considering the increase in effective sequencing depth of RPFs. Depletion with a small set of LNAs doubled the yield of RPFs in our tests, which would otherwise require twice the sequencing depth.

As illustrated in our depletion test, while a generalized set of LNAs across a multitude of conditions may not be the most effective for a specific experiment, it can substantially reduce common contaminations without the need for preliminary sequencing, thereby further reducing the barrier of entry.

The flexible nature of our approach lends itself to combination with other strategies like Ribo-FilterOut (Tomuro *et al*., 2024) or the rRNA preserving nuclease P1 (Ferguson *et al*., 2023) to obtain exceptional performance. Future endeavors should focus on expanding our initial set of contaminants by establishing well-defined and cross-validated target sets for specific organism-nuclease combinations across various tissues and conditions, thereby serving as an information resource for the entire Ribo-Seq community.

## Materials and Methods

### Biological material

All experiments were conducted using wild-type Arabidopsis thaliana (Col-0). To assess how growth conditions affect the contaminant profile, Arabidopsis plants were grown in soil for 8 days or 3 weeks, each under long-day (LD, 16 hours of light) and short-day (SD, 8 hours of light) conditions in a climate chamber at 19–22 °C. Under the same environmental conditions, seedlings were grown hydroponically for 8 days in ½-strength Murashige-Skoog (MS) medium containing 2.56 mM MES buffer, either with or without 1 % (w/v) sucrose. Etiolated seedlings were grown hydroponically in ½-strength MS medium with sucrose in total darkness after a brief period of illumination to synchronize germination. The seedlings used for testing the effectiveness of LNA depletion were grown hydroponically for 8 days in ½-strength MS medium without sucrose under short-day conditions.

### Purification of ribosomal footprints

Purification of ribosomal footprints was adapted from Hsu et al. (2016) with modifications. In short, plant material (leaf discs for 3-week-old plants, whole seedlings for 8-day-old plants) was harvested into liquid nitrogen, homogenized, and polysomes were extracted by adding 200 µl of ice-cold polysome extraction buffer (100 mM TRIS pH 8.0, 40 mM KCl, 20 mM MgCl_2_, 1 % (w/v) sodium deoxycholate or 1 % (v/v) Triton X-100, 2 % (v/v) polyoxyethylene (10) tridecyl ether, 1 mM DTT, and 355 µM (100 µg/ml) cycloheximide). The mixture was incubated on ice for 5–10 minutes and then centrifuged at 20,000 g for 7 minutes at 4 °C. The supernatant was transferred to a new tube and pooled if necessary. The total RNA concentration was determined using a Qubit 4 fluorometer with the RNA Broad Range Assay (Thermo Fisher). Footprints were generated by the addition of 0.3 U RNase If (New England Biolabs) per 1 µg of total RNA in a sample volume of 110 µl and incubation for 30 minutes at 21 °C with agitation at 300 RPM. Digested extracts were immediately loaded onto pre-equilibrated size-exclusion columns (MicroSpin S-400 HR, Cytiva) and spun for 90 seconds at 750 g to separate monosomes from free RNA fragments. The flow-through containing the monosomes was mixed with TRIzol (Invitrogen) to purify RNA. The resulting RNA pellet was resuspended in PNK reaction buffer (New England Biolabs) containing 10 mM ATP for end repair of the RNA fragments at 37 °C for 30 minutes. End-repaired RNA fragments were purified by ammonium-acetate/isopropanol precipitation and resuspended in 2x urea loading buffer. The purified RNA was separated on a denaturing 15 % polyacrylamide gel with 7 M urea (TAE buffer) at 100 V for 15 minutes and then at 150 V for 60 minutes. Bands in the size range between 18–32 nt were excised from the gel and eluted according to Reid, Shenolikar, and Nicchitta (2015). However, instead of the freeze and boil cycles, the homogenized gel pieces were sonicated for 1 hour in an ultrasonic bath. The eluted RNA was precipitated by adding one volume of isopropanol and 1.3 µl GlycoBlue coprecipitant (Invitrogen) and incubating for at least 1 hour or overnight. The precipitated RNA was washed and resuspended in 7 µl of RNase-free water.

### Library preparation and depletion of contaminants

Purified ribosomal footprints (including contaminants) were quantified with the Qubit 4 fluorometer, using the RNA High Sensitivity assay (Thermo Fisher). Between 5–20 ng of RNA was used as input for library preparation with the TrueQuant SmallRNA Seq kit (GenXPro). According to the manufacturer, when using already purified footprints at these concentrations, dilution of adapters and primers was not necessary. Libraries were prepared according to the manual with overnight incubation of the 5’-adapter, as recommended for plant RNA.

Depletion of contaminant sequences with custom LNA probes was performed during the library amplification step by adding a 10-fold LNA mix to the water volume in the PCR reaction. The LNA mix included five LNA probes (Table 1), each at a concentration of 10 µM, resulting in a final concentration of 1 µM in the master mix. Libraries were amplified with seven cycles, based on preliminary test amplifications. Final library concentrations were measured with the Qubit 4 fluorometer and the 1x DNA High Sensitivity assay (Thermo Fisher).

Sequencing was performed at GenXPro (Frankfurt, Germany), aiming for 4–7 million 75 bp single-end reads. Due to the high abundance of contaminating fragments, low sequencing depths are sufficient for preliminary testing. The trimmed reads are available at the European Nucleotide Archive (ENA) under accession number PRJEB94291.

### Data analysis and identification of contaminant sequences

Sequenced reads were quality controlled with FastQC version 0.12.1 (Andrews, 2010) and filtered with seqkit (Shen *et al*., 2016), only including reads between 20–30 nt to account for variations during gel excision. The filtered reads were aligned to the reference genome of *Arabidopsis thaliana* (TAIR 10 release) with the STAR aligner (Dobin *et al*., 2013b) using the Araport 11 Oct. 2024 release (Cheng *et al*., 2017). For accurate quantification of multimapping contaminants, we ran STAR with –outSAMmultNmax 1 and –outFilterMultimapNmax 20. Alignments in BAM format were used as input for our contamination identification script in R (version 4.0 or later). First, features were extracted from the supplied annotation file, mapped to the alignments, and counted using the packages Rsamtools (Morgan *et al*., 2024), GenomicFeatures (Lawrence *et al*., 2013), and txdbmaker (Pagès *et al*., 2024). The quantified sequences were aggregated by sorting them by their abundance, selecting the first 10,000 sequences, sorting them by length, and matching each throughout the full set of sequences (Supporting Figure S3). All matching sequences were grouped, and their summed counts as well as the shortest common sequence in the set were extracted. All matched sequences were eliminated from the search set before the following sequence was queued to increase computation speed. The final data frame, containing all aggregated sequences, was visualized using the ggplot2 package (Wickham, 2016). We generated a heatmap illustrating the distribution of similar contaminants across all samples, along with a figure of merit showing the aggregate percentage of contaminants targeted by increasing LNA set size. The contaminant identification script is available at https://github.com/ZeroG-lab/LNA-Depletion.

Features for non-multimapping reads were counted using the --quantMode GeneCounts flag in STAR. Exploratory data analysis on the counts was performed in R using the DESeq2 package (Love *et al*., 2014).

### LNA performance testing

We determined the optimal application method of the LNAs in terms of concentration and blocking step by assessing the abundance of targeted transcripts using RT-qPCR. Total RNA was extracted from leaves of 3-week-old *Arabidopsis thaliana* Col-0 wild-type plants using TRIzol reagent (Invitrogen). Purified RNA was reverse transcribed to cDNA with the iScript cDNA Synthesis Kit (Bio-Rad). The template cDNA was amplified using the PowerUP SYBR Green Master Mix in a StepOnePlus Real-Time PCR system (Applied Biosystems). The highly abundant 5.8S ribosomal RNA was chosen as a target (fwd: CGATGAAGAACGTAGCGAAA; rev: TTGTGACACCCAGGCAG). An LNA probe (CCAGGCAGACGTGCCC) targeted to the 5.8S rRNA was added in increasing concentrations at either the reverse transcription step or during the qPCR amplification. For the reverse transcription depletion, 200 ng of template RNA were used in conjunction with the LNAs. For the amplification depletions, 1 µg of template RNA was first reverse transcribed without LNAs and then 20 ng of the resulting cDNA were amplified with the LNAs. This effective 10-fold difference in template concentration between the reverse transcription and amplification steps mimics real-world workflows, where the reverse transcribed cDNA is further diluted by the PCR master mix. Target abundance was calculated from the cycle threshold differences between the tested LNA concentrations.

## Data Statement

Following the FAIR Guiding Principles, all data and code underlying this study have been made publicly available. The R script used for data processing and analysis together with detailed contaminant tables for every condition are accessible via GitHub at https://github.com/ZeroG-lab/LNA-Depletion. The next-generation sequencing data generated in this study have been submitted to the EMBL (ENA) database under accession number PRJEB94291.

## Acknowledgements

We thank Dr. Klaus Hoffmeier at GenXPro and Professor Dr. Stefan Simm at the University of Applied Sciences and Arts, Coburg, for their insightful input during the early stages of the project.

The authors declare no conflict of interest.

The research was funded by the Federal Ministry for Economic Affairs and Energy through the German Space Agency under grant numbers 50WB2130 and 50WK2270H, as well as by the European Space Agency through the ESA-CORA-GBF program. Open-access funding was enabled and organized by Projekt DEAL.

## Short Legends for Supporting Information

Supporting Figure S1: Mapping of the most abundant contaminant fragments to the structure of the 80S ribosome.

Supporting Figure S2: Comparison of RPF band intensities after electrophoretic separation with mappings to transcript origin along the fragment length.

Supporting Figure S3: Flow of the contaminant identification script.

Supporting Tables: Shortest common sequences of the top 30 contaminant fragments in *Arabidopsis thaliana* for each tested growth condition.

## Author Contributions

CRediT: Conceptualization: DAR, FEP, MB; Data curation: DAR; Formal Analysis: DAR, FEP, MB; Funding acquisition: MB; Investigation: DAR, FEP; Methodology: DAR, FEP, MB; Project administration: MB; Resources: MB; Software: DAR, MB; Visualization: DAR, FEP; Writing – original draft: DAR; Writing – review & editing: DAR, MB

