## Supporting Information Legends for "Data-driven design of LNA-blockers for efficient contaminant removal in Ribo-seq libraries"

**Supporting Tables S1-S9:** Legends included in Supporting_Tables.docx.

**Supporting Figure S1:** Mapping the most abundant contaminating rRNA fragments from previous Ribo-Seq experiments onto the structure of the 80S ribosome from tobacco. The ribosome is shown with the large 60S subunit in different shades of beige and the small 40S subunit in shades of purple, highlighting their structural components. Within these subunits, the rRNA sequences are shaded in matching colors, with the 5.8S rRNA distinctly highlighted in green. Contaminating rRNA fragments detected in Ribo-Seq libraries are marked red on their respective rRNA sequences. To provide a comprehensive view, the ribosome is rotated 180 degrees, as indicated by arrows, revealing additional contaminating fragments. Specific fragments are shown in a zoomed-in perspective, with annotations detailing their originating rRNA sequences provided below the close-ups. The structure of the 80S ribosome is based on a Cryo-EM model solved by Smirnova et al. (2023) and visualized using PyMOL (The PyMOL Molecular Graphics System, Version 3.0 Schrödinger, LLC.).

**Supporting Figure S2:** Identification of RNA classes within the size range of ribosome-protected footprints (RPFs). The gel image on the left shows control samples not treated with RNase I (indicated by "-") and samples treated with RNase I (indicated by "+"), where distinct bands appear due to nuclease digestion. The middle panel presents the intensity profile of these bands, measured using ImageJ. On the right, the sequencing read distribution profile of the nuclease-digested samples exhibits a high resemblance to the gel intensity profile, confirming that the visible bands mainly consist of non-coding RNA fragments rather than RPFs.

**Supporting Figure S3:** Flow of the contaminant identification script**.** Alignments in BAM format from a small-scale preliminary sequencing experiment are supplied to our custom R-script as input. Iterative aggregation of identified sequences produces a grouped output of contaminants that are suited as targets for LNA-based depletion. The results are visualized in a heatmap and figure of merit, as presented in figure 1. Data sets are colored blue, functions are colored green.
