## Supporting Tables S1-9 for "Data-driven design of LNA-blockers for efficient contaminant removal in Ribo-seq libraries"

Supporting Table S1: Shortest common sequences of most abundant fragment groups for Arabidopsis thaliana grown for **3 weeks on soil under long-day conditions**. Mean percentage is the relation of fragments sharing the indicated sequence to the total sum of all reads. Additional Gene IDs and biotype information are available in the extended tables in the corresponding GitHub repository.

| Sequence | Mean Percentage | Cumulative Percentage |
| --- | --- | --- |
| ACGTCTGCCTGGGTGTCACA | 6.9 | 6.9 |
| GGCCGGGGGACGGACTGGGA | 5.32 | 12.22 |
| CTGTTTTTGGTCCCAAGGCT | 3.25 | 15.48 |
| CGTCGCTGCCGTGATCGTGG | 1.67 | 17.14 |
| GGTCGGCCAGTGAGACGGTG | 1.64 | 18.78 |
| CCACTTGACACCTATCGTAA | 1.54 | 20.32 |
| TTACGATAGGTGTCAAGTGG | 1.51 | 21.84 |
| GCTCTCCCACAACCCCGTTT | 1.4 | 23.24 |
| AAACGGGGTTGTGGGAGAGC | 1.4 | 24.64 |
| TGGATACTAGGCGCTGTGCG | 1.36 | 26 |
| CGCACAGCGCCTAGTATCCA | 1.36 | 27.35 |
| AACTCTGAATACTAGATATG | 1.28 | 28.63 |
| CTGATGATTCATGATAACTC | 1.26 | 29.89 |
| CGAGGGCACGTCTGCCTGGG | 1.16 | 31.05 |
| AGGTCATATCTAGTATTCAG | 1.01 | 32.06 |
| TGAATACTAGATATGACCTC | 0.95 | 33.02 |
| GGCAGTCTGTTCAGGGTTCCA | 0.92 | 33.93 |
| TGGAACCCTGAACAGACTGCC | 0.91 | 34.84 |
| GCTCGTCTCGCCGTGACCTTC | 0.88 | 35.73 |
| GAAGGTCACGGCGAGACGAGC | 0.88 | 36.61 |
| CGCGCTACACTGATGTATTCA | 0.85 | 37.46 |
| GCGGGTGCACCGCCGACCGAC | 0.77 | 38.23 |
| CCCAGGTGGCGAAGAGCCGAC | 0.63 | 38.86 |
| GTCGGCTCTTCGCCACCTGGG | 0.63 | 39.5 |
| AGAGAGTTGTCTCGCGCCCCTA | 0.63 | 40.13 |
| TAGGGGCGCGAGACAACTCTCT | 0.63 | 40.76 |
| AAATCTCAACCAGCCACTGC | 0.6 | 41.36 |
| TAACCCATGCTATACTCCCA | 0.6 | 41.96 |
| CTGAATACTAGATATGACCT | 0.59 | 42.55 |
| TGGGAGTATAGCATGGGTTA | 0.58 | 43.14 |

Supporting Table S2: Shortest common sequences of most abundant fragment groups for Arabidopsis thaliana grown for **3 weeks on soil under short-day conditions**. Mean percentage is the relation of fragments sharing the indicated sequence to the total sum of all reads. Additional Gene IDs and biotype information are available in the extended tables in the corresponding GitHub repository.

| Sequence | Mean Percentage | Cumulative Percentage |
| --- | --- | --- |
| GGCCGGGGGACGGACTGGGA | 7.98 | 7.98 |
| AAACGGGGTTGTGGGAGAGC | 3.9 | 11.88 |
| GCTCTCCCACAACCCCGTTT | 3.89 | 15.77 |
| ACGTCTGCCTGGGTGTCACA | 3.43 | 19.2 |
| ACGCATTTATTAGATAAAAG | 2.99 | 22.19 |
| CGGCTTGTCCCTTCGGTCGG | 1.9 | 24.09 |
| CTGTTTTTGGTCCCAAGGCT | 1.64 | 25.73 |
| GAAGGTCACGGCGAGACGAGC | 1.48 | 27.21 |
| GCTCGTCTCGCCGTGACCTTC | 1.47 | 28.68 |
| CGCGCTACACTGATGTATTCA | 1.21 | 29.89 |
| GTTGGTGGAGCGATTTGTCT | 1.13 | 31.02 |
| CGAGGGCACGTCTGCCTGGG | 1.09 | 32.11 |
| GCGAGCTACCAAACTGCTCTACCCCG | 1.08 | 33.19 |
| TGCCGGCCGGGGGACGGACT | 0.96 | 34.15 |
| GATGCCGACCAGGGATCAGC | 0.87 | 35.02 |
| CTGATGATTCATGATAACTC | 0.85 | 35.87 |
| CGCACAGCGCCTAGTATCCA | 0.79 | 36.66 |
| TGGATACTAGGCGCTGTGCG | 0.78 | 37.44 |
| GATATCCTAACCGCTGGACGACATCGGA | 0.72 | 38.16 |
| TGGTCGGCTTGTCCCTTCGG | 0.7 | 38.86 |
| CCGACCTGCCGGAAACGAACCAGCGAC | 0.63 | 39.48 |
| CGTCGCTGCCGTGATCGTGG | 0.58 | 40.06 |
| AGCCCAGCTTGAAAATCGGA | 0.51 | 40.57 |
| AAATCTCAACCAGCCACTGC | 0.49 | 41.07 |
| GCGGGTGCACCGCCGACCGAC | 0.49 | 41.55 |
| TAACCGCTGGACGATAACGGA | 0.47 | 42.02 |
| GGTCGGCCAGTGAGACGGTG | 0.43 | 42.45 |
| GTCGCTGGTTCGTTTCCGGCAGGTCGG | 0.43 | 42.89 |
| ATGATTAACAGGGACAGTCGGG | 0.43 | 43.31 |
| AAAATAGCTCGACGCCAGGA | 0.4 | 43.71 |

Supporting Table S3: Shortest common sequences of most abundant fragment groups for Arabidopsis thaliana grown for **8 days on soil under long-day conditions**. Mean percentage is the relation of fragments sharing the indicated sequence to the total sum of all reads. Additional Gene IDs and biotype information are available in the extended tables in the corresponding GitHub repository.

| Sequence | Mean Percentage | Cumulative Percentage |
| --- | --- | --- |
| GGCCGGGGGACGGACTGGGA | 8.04 | 8.04 |
| AAACGGGGTTGTGGGAGAGC | 5.51 | 13.55 |
| GCTCTCCCACAACCCCGTTT | 5.47 | 19.02 |
| CGGCTTGTCCCTTCGGTCGG | 3.35 | 22.37 |
| ACGCATTTATTAGATAAAAG | 3.08 | 25.46 |
| TGGTCGGCTTGTCCCTTCGG | 1.82 | 27.28 |
| CTGTTTTTGGTCCCAAGGCT | 1.55 | 28.83 |
| ACGTCTGCCTGGGTGTCACA | 1.54 | 30.36 |
| CTGATGATTCATGATAACTC | 1.05 | 31.42 |
| CGCACAGCGCCTAGTATCCA | 0.85 | 32.27 |
| TGGATACTAGGCGCTGTGCG | 0.85 | 33.12 |
| CCGACCTGCCGGAAACGAACCAGCGAC | 0.82 | 33.94 |
| TGCCGGCCGGGGGACGGACT | 0.78 | 34.71 |
| GGCAGTCTGTTCAGGGTTCCA | 0.76 | 35.47 |
| TGGAACCCTGAACAGACTGCC | 0.75 | 36.22 |
| GATGCCGACCAGGGATCAGC | 0.74 | 36.96 |
| GTTGGTGGAGCGATTTGTCT | 0.7 | 37.65 |
| AAAACGGGGTTGTGGGAGAG | 0.68 | 38.33 |
| CTCTCCCACAACCCCGTTTT | 0.68 | 39.01 |
| GCTCGTCTCGCCGTGACCTTC | 0.67 | 39.67 |
| GAAGGTCACGGCGAGACGAGC | 0.65 | 40.33 |
| CGAGGGCACGTCTGCCTGGG | 0.6 | 40.93 |
| ATGATTAACAGGGACAGTCGGG | 0.57 | 41.5 |
| GTCGCTGGTTCGTTTCCGGCAGGTCGG | 0.57 | 42.07 |
| GCGAGCTACCAAACTGCTCTACCCCG | 0.57 | 42.64 |
| CCCACCACGCTTCCGCTGCGCCACTC | 0.54 | 43.18 |
| CGCGCTACACTGATGTATTCA | 0.47 | 43.65 |
| ACATAGTAAGGATTGACAGA | 0.44 | 44.08 |
| GGGGACGGACTGGGAACGGCT | 0.43 | 44.52 |
| AAATCTCAACCAGCCACTGC | 0.41 | 44.93 |

Supporting Table S4: Shortest common sequences of most abundant fragment groups for Arabidopsis thaliana grown for **8 days on soil under short-day conditions**. Mean percentage is the relation of fragments sharing the indicated sequence to the total sum of all reads. Additional Gene IDs and biotype information are available in the extended tables in the corresponding GitHub repository.

| Sequence | Mean Percentage | Cumulative Percentage |
| --- | --- | --- |
| GGCCGGGGGACGGACTGGGA | 6.95 | 6.95 |
| CGGCTTGTCCCTTCGGTCGG | 5.51 | 12.45 |
| ACGCATTTATTAGATAAAAG | 5.3 | 17.75 |
| AAACGGGGTTGTGGGAGAGC | 3.62 | 21.37 |
| GCTCTCCCACAACCCCGTTT | 3.61 | 24.98 |
| TGGTCGGCTTGTCCCTTCGG | 2.89 | 27.86 |
| ACGTCTGCCTGGGTGTCACA | 1.92 | 29.78 |
| CTGTTTTTGGTCCCAAGGCT | 1.89 | 31.67 |
| CGAGGGCACGTCTGCCTGGG | 1.17 | 32.84 |
| CTGATGATTCATGATAACTC | 1.13 | 33.97 |
| GATGCCGACCAGGGATCAGC | 1 | 34.97 |
| GCTCGTCTCGCCGTGACCTTC | 0.97 | 35.95 |
| GAAGGTCACGGCGAGACGAGC | 0.97 | 36.91 |
| TGCCGGCCGGGGGACGGACT | 0.75 | 37.66 |
| AGCCCAGCTTGAAAATCGGA | 0.66 | 38.33 |
| CGCACAGCGCCTAGTATCCA | 0.66 | 38.99 |
| TGGATACTAGGCGCTGTGCG | 0.65 | 39.64 |
| GTTGGTGGAGCGATTTGTCT | 0.62 | 40.26 |
| AAATCTCAACCAGCCACTGC | 0.6 | 40.87 |
| CTACGCCTAGGACACCAGAA | 0.54 | 41.41 |
| CCGACCTGCCGGAAACGAACCAGCGAC | 0.51 | 41.92 |
| GCCGCTCACGCCCGGTCGTA | 0.46 | 42.37 |
| CGCGCTACACTGATGTATTCA | 0.45 | 42.82 |
| CTCTCCCACAACCCCGTTTT | 0.39 | 43.21 |
| AAAACGGGGTTGTGGGAGAG | 0.38 | 43.59 |
| GTCGCTGGTTCGTTTCCGGCAGGTCGG | 0.34 | 43.93 |
| CCCACCACGCTTCCGCTGCGCCACTC | 0.34 | 44.28 |
| GATATCCTAACCGCTGGACGACATCGGA | 0.34 | 44.61 |
| GGCAGTCTGTTCAGGGTTCCA | 0.32 | 44.93 |
| TGGAACCCTGAACAGACTGCC | 0.32 | 45.25 |

Supporting Table S5: Shortest common sequences of most abundant fragment groups for Arabidopsis thaliana grown **hydroponically (Murashige and Skoog medium + sucrose) for 8 days in the dark (etiolated)**. Mean percentage is the relation of fragments sharing the indicated sequence to the total sum of all reads. Additional Gene IDs and biotype information are available in the extended tables in the corresponding GitHub repository.

| Sequence | Mean Percentage | Cumulative Percentage |
| --- | --- | --- |
| ACGTCTGCCTGGGTGTCACA | 18.18 | 18.18 |
| CTGTTTTTGGTCCCAAGGCT | 9.49 | 27.67 |
| CGAGGGCACGTCTGCCTGGG | 4.12 | 31.78 |
| CGTCGCTGCCGTGATCGTGG | 3.2 | 34.98 |
| CGAGGCAAACTCTGAATACT | 2.21 | 37.19 |
| CTGATGATTCATGATAACTC | 1.93 | 39.12 |
| GCGGGTGCACCGCCGACCGAC | 1.9 | 41.02 |
| GGCCGGGGGACGGACTGGGA | 1.56 | 42.57 |
| GGCGTTCGAATTGTAGTCTG | 1.52 | 44.1 |
| GATGCCGACCAGGGATCAGC | 1.48 | 45.57 |
| CCGACCCTCAGTAGGAGCTT | 1.47 | 47.05 |
| CCGACGCGGATTACGGTGGCG | 1.47 | 48.52 |
| AGTATTCAGAGTTTGCCTCG | 1.32 | 49.84 |
| GGACCGACGCGGATTACGGT | 1.1 | 50.93 |
| CGCGGATTACGGTGGCGGCCTA | 0.84 | 51.78 |
| ATGATTAACAGGGACAGTCGGG | 0.79 | 52.56 |
| CGCGCTACACTGATGTATTCA | 0.65 | 53.22 |
| CCGGAGGTAGGGTCCAGCGG | 0.51 | 53.73 |
| CCCCTGTTTTTGGTCCCAAG | 0.5 | 54.23 |
| GCCGCTCACGCCCGGTCGTA | 0.4 | 54.63 |
| AAATCTCAACCAGCCACTGC | 0.38 | 55.01 |
| CGGTGAAGTGTTCGGATCGCG | 0.35 | 55.36 |
| CGGGCTAGAAGCGACGCATG | 0.33 | 55.68 |
| ATGATAACTCGACGGATCGC | 0.33 | 56.01 |
| CGCGAGAAGTCCACTAAACC | 0.29 | 56.31 |
| TGTTTTTGGTCCCAAGGCTC | 0.29 | 56.6 |
| ACAGCCTGCCCACCCTGGAA | 0.28 | 56.88 |
| AGCCCAGCTTGAAAATCGGA | 0.27 | 57.15 |
| AGCCCTTTGTCGCTAAGATTCGA | 0.27 | 57.42 |
| GGTGCACCGCCGACCGACCTTG | 0.25 | 57.67 |

Supporting Table S6: Shortest common sequences of most abundant fragment groups for Arabidopsis thaliana grown **hydroponically (Murashige and Skoog medium without sucrose) for 8 days under long-day conditions**. Mean percentage is the relation of fragments sharing the indicated sequence to the total sum of all reads. Additional Gene IDs and biotype information are available in the extended tables in the corresponding GitHub repository.

| Sequence | Mean Percentage | Cumulative Percentage |
| --- | --- | --- |
| GGCCGGGGGACGGACTGGGA | 13.37 | 13.37 |
| ACGTCTGCCTGGGTGTCACA | 8.14 | 21.51 |
| CGGCTTGTCCCTTCGGTCGG | 4.76 | 26.27 |
| CTGTTTTTGGTCCCAAGGCT | 2.58 | 28.85 |
| AAACGGGGTTGTGGGAGAGC | 2.28 | 31.13 |
| GCTCTCCCACAACCCCGTTT | 2.28 | 33.41 |
| ACGCATTTATTAGATAAAAG | 2.25 | 35.65 |
| CGAGGGCACGTCTGCCTGGG | 1.99 | 37.65 |
| TGGATACTAGGCGCTGTGCG | 1.57 | 39.21 |
| CGCACAGCGCCTAGTATCCA | 1.56 | 40.78 |
| TGCCGGCCGGGGGACGGACT | 1.44 | 42.22 |
| CTGATGATTCATGATAACTC | 1.43 | 43.66 |
| GATGCCGACCAGGGATCAGC | 1.36 | 45.01 |
| CGAGGCAAACTCTGAATACT | 1.34 | 46.36 |
| TGGTCGGCTTGTCCCTTCGG | 1.33 | 47.69 |
| GCGGGTGCACCGCCGACCGAC | 0.92 | 48.61 |
| AGTATTCAGAGTTTGCCTCG | 0.82 | 49.43 |
| CGCGCTACACTGATGTATTCA | 0.68 | 50.11 |
| AGCCCAGCTTGAAAATCGGA | 0.63 | 50.74 |
| GAAGGTCACGGCGAGACGAGC | 0.62 | 51.36 |
| GCTCGTCTCGCCGTGACCTTC | 0.62 | 51.98 |
| CGTCGCTGCCGTGATCGTGG | 0.62 | 52.6 |
| TGGAAGGGACGCATTTATTA | 0.56 | 53.16 |
| CCGACCCTCAGTAGGAGCTT | 0.51 | 53.67 |
| CCGGAGGTAGGGTCCAGCGG | 0.49 | 54.16 |
| ATGATTAACAGGGACAGTCGGG | 0.46 | 54.62 |
| GCCGCTCACGCCCGGTCGTA | 0.45 | 55.07 |
| CCACTTGACACCTATCGTAA | 0.45 | 55.52 |
| TTACGATAGGTGTCAAGTGG | 0.44 | 55.96 |
| GTTGGTGGAGCGATTTGTCT | 0.43 | 56.4 |

Supporting Table S7: Shortest common sequences of most abundant fragment groups for Arabidopsis thaliana grown **hydroponically (Murashige and Skoog medium with sucrose) for 8 days under long-day conditions**. Mean percentage is the relation of fragments sharing the indicated sequence to the total sum of all reads. Additional Gene IDs and biotype information are available in the extended tables in the corresponding GitHub repository.

| Sequence | Mean Percentage | Cumulative Percentage |
| --- | --- | --- |
| GGCCGGGGGACGGACTGGGA | 13.19 | 13.19 |
| CGGCTTGTCCCTTCGGTCGG | 5.91 | 19.11 |
| ACGTCTGCCTGGGTGTCACA | 4.67 | 23.78 |
| CTGTTTTTGGTCCCAAGGCT | 2.92 | 26.7 |
| ACGCATTTATTAGATAAAAG | 2.53 | 29.23 |
| TGGAAGGGACGCATTTATTA | 2.36 | 31.6 |
| CGAGGCAAACTCTGAATACT | 2.09 | 33.68 |
| ATGATTAACAGGGACAGTCGGG | 1.93 | 35.62 |
| GGCGTTCGAATTGTAGTCTG | 1.83 | 37.45 |
| TGGTCGGCTTGTCCCTTCGG | 1.78 | 39.23 |
| CGAGGGCACGTCTGCCTGGG | 1.72 | 40.94 |
| GATGCCGACCAGGGATCAGC | 1.69 | 42.64 |
| TGCCGGCCGGGGGACGGACT | 1.5 | 44.13 |
| AGTATTCAGAGTTTGCCTCG | 1.25 | 45.39 |
| TGGATACTAGGCGCTGTGCG | 1.25 | 46.63 |
| CGCACAGCGCCTAGTATCCA | 1.24 | 47.88 |
| GCTCGTCTCGCCGTGACCTTC | 1.08 | 48.96 |
| GAAGGTCACGGCGAGACGAGC | 1.08 | 50.04 |
| GCTCTCCCACAACCCCGTTT | 1 | 51.04 |
| AAACGGGGTTGTGGGAGAGC | 0.99 | 52.03 |
| CGTCGCTGCCGTGATCGTGG | 0.74 | 52.77 |
| CCGGAGGTAGGGTCCAGCGG | 0.68 | 53.45 |
| CTGATGATTCATGATAACTC | 0.67 | 54.12 |
| AGCCCAGCTTGAAAATCGGA | 0.67 | 54.79 |
| AAATCTCAACCAGCCACTGC | 0.59 | 55.38 |
| CGCGCTACACTGATGTATTCA | 0.55 | 55.93 |
| GTTGGTGGAGCGATTTGTCT | 0.54 | 56.47 |
| GCGGGTGCACCGCCGACCGAC | 0.49 | 56.96 |
| CGGCCGGGGGACGGACTGGG | 0.39 | 57.35 |
| GCGTTCGAATTGTAGTCTGG | 0.37 | 57.72 |

Supporting Table S8: Shortest common sequences of most abundant fragment groups for Arabidopsis thaliana grown **hydroponically (Murashige and Skoog medium without sucrose) for 8 days under short-day conditions**. Mean percentage is the relation of fragments sharing the indicated sequence to the total sum of all reads. Additional Gene IDs and biotype information are available in the extended tables in the corresponding GitHub repository.

| Sequence | Mean Percentage | Cumulative Percentage |
| --- | --- | --- |
| ACGTCTGCCTGGGTGTCACA | 9.76 | 9.76 |
| GGCCGGGGGACGGACTGGGA | 6.93 | 16.69 |
| CGGCTTGTCCCTTCGGTCGG | 3.67 | 20.36 |
| CTGTTTTTGGTCCCAAGGCT | 3.18 | 23.54 |
| CTGATGATTCATGATAACTC | 2.74 | 26.28 |
| ACGCATTTATTAGATAAAAG | 2.58 | 28.86 |
| CGAGGGCACGTCTGCCTGGG | 2.46 | 31.32 |
| CGAGGCAAACTCTGAATACT | 1.34 | 32.66 |
| TGGATACTAGGCGCTGTGCG | 1.3 | 33.96 |
| CGCACAGCGCCTAGTATCCA | 1.3 | 35.26 |
| GATGCCGACCAGGGATCAGC | 1.25 | 36.5 |
| GCTCTCCCACAACCCCGTTT | 1.21 | 37.71 |
| AAACGGGGTTGTGGGAGAGC | 1.21 | 38.91 |
| TGCCGGCCGGGGGACGGACT | 1.12 | 40.03 |
| GCGGGTGCACCGCCGACCGAC | 0.96 | 40.99 |
| TGGTCGGCTTGTCCCTTCGG | 0.91 | 41.9 |
| AGTATTCAGAGTTTGCCTCG | 0.8 | 42.7 |
| CGTCGCTGCCGTGATCGTGG | 0.73 | 43.43 |
| GAAGGTCACGGCGAGACGAGC | 0.72 | 44.15 |
| GCTCGTCTCGCCGTGACCTTC | 0.71 | 44.85 |
| CGCGCTACACTGATGTATTCA | 0.69 | 45.54 |
| ATGATTAACAGGGACAGTCGGG | 0.59 | 46.13 |
| TGGAAGGGACGCATTTATTA | 0.58 | 46.72 |
| CCGACCCTCAGTAGGAGCTT | 0.52 | 47.24 |
| CCACTTGACACCTATCGTAA | 0.52 | 47.76 |
| TTACGATAGGTGTCAAGTGG | 0.52 | 48.27 |
| GCCGCTCACGCCCGGTCGTA | 0.48 | 48.75 |
| AGCCCAGCTTGAAAATCGGA | 0.46 | 49.21 |
| AAAATAGCTCGACGCCAGGA | 0.45 | 49.66 |
| TCCTGGCGTCGAGCTATTTT | 0.44 | 50.1 |

Supporting Table S9: Shortest common sequences of most abundant fragment groups for Arabidopsis thaliana grown **hydroponically (Murashige and Skoog medium with sucrose) for 8 days under short-day conditions**. Mean percentage is the relation of fragments sharing the indicated sequence to the total sum of all reads. Additional Gene IDs and biotype information are available in the extended tables in the corresponding GitHub repository.

| Sequence | Mean Percentage | Cumulative Percentage |
| --- | --- | --- |
| ACGTCTGCCTGGGTGTCACA | 12.52 | 12.52 |
| GGCCGGGGGACGGACTGGGA | 7.04 | 19.56 |
| CTGTTTTTGGTCCCAAGGCT | 5.36 | 24.92 |
| CGGCTTGTCCCTTCGGTCGG | 4.12 | 29.04 |
| ACGCATTTATTAGATAAAAG | 2.86 | 31.91 |
| CGAGGGCACGTCTGCCTGGG | 2.64 | 34.54 |
| GATGCCGACCAGGGATCAGC | 1.9 | 36.44 |
| CTGATGATTCATGATAACTC | 1.7 | 38.15 |
| TGGAAGGGACGCATTTATTA | 1.64 | 39.79 |
| GCGGGTGCACCGCCGACCGAC | 1.63 | 41.42 |
| TGGTCGGCTTGTCCCTTCGG | 1.14 | 42.56 |
| CGAGGCAAACTCTGAATACT | 1.12 | 43.68 |
| CGTCGCTGCCGTGATCGTGG | 1.05 | 44.73 |
| GGCGTTCGAATTGTAGTCTG | 0.97 | 45.71 |
| TGCCGGCCGGGGGACGGACT | 0.83 | 46.54 |
| CGCGCTACACTGATGTATTCA | 0.81 | 47.35 |
| GCCGACCCTCAGTAGGAGCT | 0.76 | 48.11 |
| CCGACCCTCAGTAGGAGCTT | 0.71 | 48.82 |
| GCTCTCCCACAACCCCGTTT | 0.7 | 49.52 |
| AAACGGGGTTGTGGGAGAGC | 0.69 | 50.21 |
| TGGATACTAGGCGCTGTGCG | 0.68 | 50.89 |
| CGCACAGCGCCTAGTATCCA | 0.68 | 51.57 |
| AGTATTCAGAGTTTGCCTCG | 0.66 | 52.23 |
| ATGATTAACAGGGACAGTCGGG | 0.64 | 52.87 |
| CCCCTTGGGGTTGTGGGACG | 0.56 | 53.43 |
| GAAGGTCACGGCGAGACGAGC | 0.48 | 53.9 |
| GCTCGTCTCGCCGTGACCTTC | 0.48 | 54.38 |
| CCGGAGGTAGGGTCCAGCGG | 0.34 | 54.72 |
| GTTGGTGGAGCGATTTGTCT | 0.34 | 55.06 |
| GGACCGACGCGGATTACGGT | 0.34 | 55.4 |
