## Supplementary figures and images for "Data-driven design of LNA-blockers for efficient contaminant removal in Ribo-seq libraries"

### Supporting Figure S1

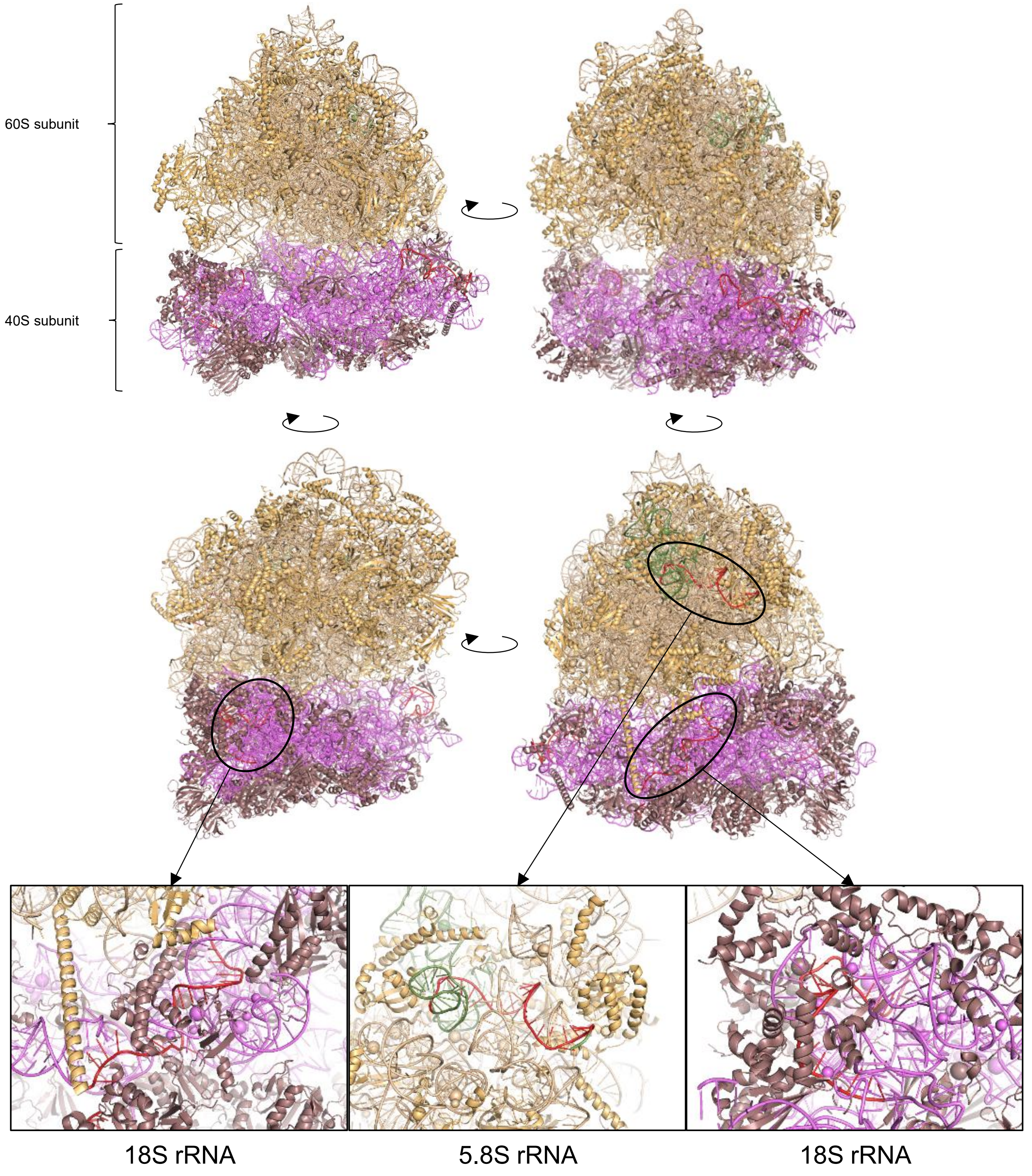

### Supporting Figure S2

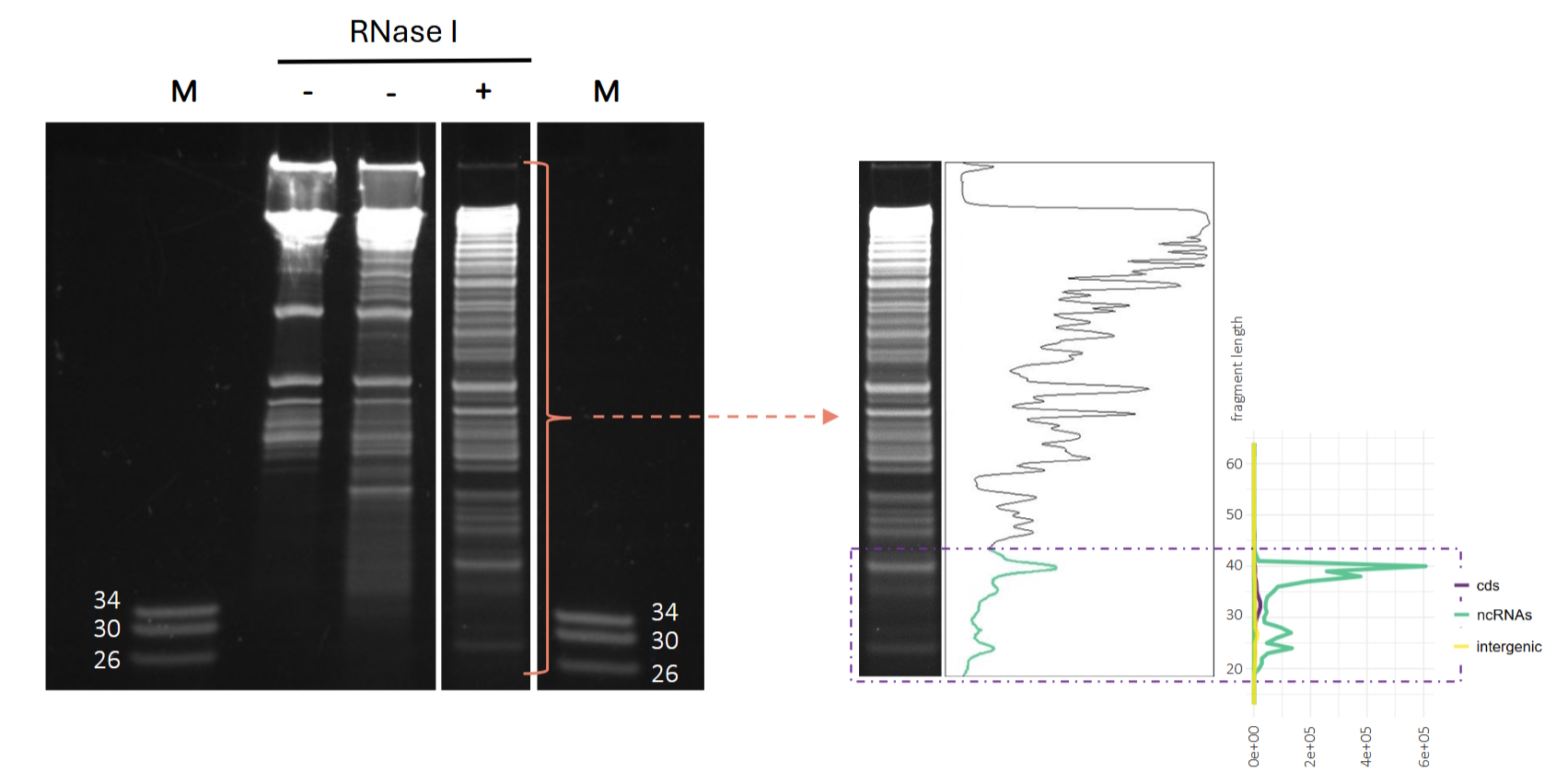

### Supporting Figure S3

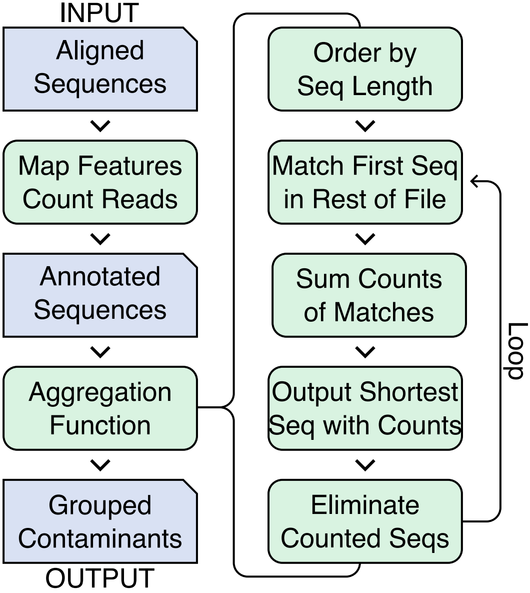
